# Descriptive pan-cancer genetic analysis of disulfidptosis-related gene set

**DOI:** 10.1101/2023.02.25.529997

**Authors:** Hengrui Liu, Tao Tang

## Abstract

**Background:** A recent study has identified a novel programmed cell death pathway, termed disulfidoptosis, which is based on disulfide proteins. This discovery provides new insight into the mechanisms of cell death and may have implications for therapeutic strategies targeting cell death pathways. This study aimed to evaluate the pan-cancer genomics and clinical association of disulfidptosis and disulfidptosis-related cell death genes, including SLC7A11, NADPH, INF2, CD2AP, PDLIM1, ACTN4, MYH9, MYH10, IQGAP1, FLNA, FLNB, TLN1, MYL6, ACTB, DSTN, and CAPZB.

**Methods:** Using multi-omics profiling data, this study provides a comprehensive and systematic characterization of disulfidptosis genes across more than 9000 samples of over 30 types of cancer.

**Results:** FLNA and FLNB were the two most frequently mutated disulfidptosis cell death genes in cancer. UCEC and SKCM were the two cancer types that have the highest mutation rates while the mutation of ACTN4 was associated with worse survival of CESC and ESCA. Breast cancer was potentially affected by disulfidptosis because its subtypes are different in disulfidptosis gene expression. Similarly, KIRC might also be associated with disulfidptosis.). Additionally, the association of disulfidptosis-related cell death genes with survival was analyzed, with MESO and LGG as the top cancer types with survival associated with disulfidptosis cell death genes. The correlation between CNV and survival across multiple cancer types found that UCEC, KIRP, LGG, and KIRC were the top cancer types where the CNV level was associated with survival. There was a negative correlation between expression and methylation for most of the genes and there was only a slight correlation between methylation levels and survival of cancer in LGG. About half of the disulfidptosis-related cell death proteins were associated with the activation of EMT. Disulfidptosis genes were correlated to immune cell infiltration levels in cancers. Multiple compounds were identified as potential drugs that might be affected by disulfidptosis-related cell death for future study.

**Conclusion:** Disulfidptosis cell death genes are potentially involved in many cancer types and can be developed as candidates for cancer diagnosis, prognosis, and therapeutic biomarkers.

## Introduction

In recent years, cancer has become a major public health concern across the world, with a projected 1,918,030 new cancer cases and 609,360 cancer deaths in the United States alone in 2022 according to the American Cancer Society[1]. Cancer is a complex and multifaceted disease that requires comprehensive research in order to improve early detection and treatment methods. By doing so, it is possible to reduce the number of cancer-related deaths and help more individuals to achieve successful treatment outcomes.

The fast-growing studies of programmed cell death have boosted a perspective on its usage in cancer diagnosis, prognosis, and therapeutics[2-4]. A recent paper[5] has revealed another programmed cell death, the disulfidptosis, a programmed cell death based on disulfide proteins. The paper describes a previously uncharacterized form of cell death, termed disulfidptosis, mediated by aberrant intracellular disulfide accumulation in cystine transporter solute carrier family 7 member 11 (SLC7A11)-high cells under glucose starvation. This cell death is distinct from apoptosis and ferroptosis, and is triggered by the susceptibility of the actin cytoskeleton to disulfide stress. Chemical proteomics and cell biological analyses revealed that glucose starvation in SLC7A11high cells induces aberrant disulfide bonds in actin cytoskeleton proteins, leading to F-actin collapse in a SLC7A11-dependent manner. CRISPR screens and functional studies further revealed that inactivation of the WAVE regulatory complex suppresses disulfidptosis, whereas constitutive activation of Rac promotes disulfidptosis. Additionally, glucose transporter inhibitors were found to induce disulfidptosis in SLC7A11high cancer cells, which in turn suppressed SLC7A11high tumour growth.

In order to better understand the molecular mechanisms underlying cancer development and progression, it is essential to study the genetic and molecular alterations of different cancer types. To this end, cancer databases, such as The Cancer Genome Atlas (TCGA) [6], offer a wealth of information, providing gene alteration, gene expression, and clinical survival data on different cancer types. This allows for the undertaking of pan-cancer studies to identify and gain insights into potential biomarkers and therapeutic targets for cancer treatments. By leveraging the power of such databases, researchers can better understand the complexity of cancer and develop more effective treatments and prevention strategies[7-9].

This study focused on 16 genes in disulfidptosis-related cell death, including SLC7A11, NADPH, INF2, CD2AP, PDLIM1, ACTN4, MYH9, MYH10, IQGAP1, FLNA, FLNB, TLN1, MYL6, ACTB, DSTN, and CAPZB. SLC7A11, as above mentioned, is the core molecule for disulfidptosis. A previous pan-cancer analysis revealed that the SLC7A11 gene is a clinical prognostic biomarker for Cancer. Disulfidptosis is mediated by NADPH, hence, this gene is also critical to be observed for disulfidptosis in cancer. INF2, CD2AP, PDLIM1, ACTN4, MYH9, MYH10, IQGAP1, FLNA, FLNB, TLN1, MYL6, ACTB, DSTN, and CAPZB were the genes encoding actin-related proteins with disulfide bonds increased following the glucose starvation [5]. The formation of disulfide bonds in actin cytoskeleton proteins due to glucose starvation has been proposed to be a consequence of NADPH depletion, rather than a secondary consequence of cell death. This phenomenon has been studied extensively in the context of cancer, where glucose deprivation is a common feature of the tumour microenvironment. Therefore, these actin cytoskeleton proteins are very likely to mediate disulfidptosis.

This study aimed to provide a comprehensive pan-cancer genomics and clinical association profile of these disulfidptosis cell death genes for future reference. We think, with the rise of disulfidptosis cancer research, these in-time profiles will provide a genetic overview and useful information for future studies on the role of disulfidptosis-related cell death in cancers.

## Methods

### 1. Data acquisitions

For the purpose of this study, data was obtained from a variety of sources including The Cancer Genome Atlas (TCGA) [6], NCI Genomic Data Commons (TCGA), the Cancer Proteome Atlas (TCPA), numerous databases containing experimentally verified and predicted miRNA regulation data (TarBase, miRTarBase, mir2disease, Targetscan, and miRanda), the Tumor Immune Dysfunction and Exclusion (TIDE) database, and the Drug Sensitivity in Cancer (GDSC) and Cancer Therapeutics Response Portal (CTRP) databases. Single-nucleotide variants (SNVs) and copy number variants (CNVs) were sourced from the NCI Genomic Data Commons (TCGA), methylation data was obtained from TCGA, and reverse-phase protein array (RPPA) data was attained from the Cancer Proteome Atlas (TCPA). Additionally, gene-drug sensitivity data was downloaded from both the Drug Sensitivity in Cancer (GDSC) database and the Cancer Therapeutics Response Portal (CTRP).

### 2. Gene alterations and expression analysis

The analysis of expression and methylation were conducted using R Foundation for Statistical Computing (version 4.0.3) and the ggplot2 (version 3.3.2). SNP plots were generated by the maftools[22], and CNV data were processed with GISTICS2.0 [23].

### 3. Pathway activity analysis

The Reverse-phase Protein Array (RPPA) data from the TCPA database was used to calculate pathway scores for 7876 samples. Ten cancer-related pathways were included, such as the tuberous sclerosis 1 protein (TSC)/mechanistic target of rapamycin (mTOR) and Phosphoinositide 3-kinases (PI3K)/protein kinase B (AKT) pathways. The pathway score was determined by summing the relative protein level of all positive regulatory components and subtracting the levels of negative regulatory components. The pathway activity score (PAS) was estimated using a method analogous to that used in previous studies[10,11]; gene expression was divided into two groups (High and Low) based on the median expression, and the difference in PAS between the two groups was analyzed using a Student′s t-test. Significance was determined by adjusting the P-value using the False Discovery Rate (FDR) method, with an FDR ≤0.05 indicating significance. If the PAS (Gene A group High) was greater than the PAS (Gene A group Low), then gene A was determined to have an activating effect on the pathway; otherwise, it had an inhibitory effect.

### 4. MicroRNA (miRNA) regulation network analysis

The correlation between miRNA and gene expression in cancer samples was evaluated using the Pearson product-moment correlation coefficient and the t-distribution. Specifically, miRNA and gene expression data from the Cancer Genome Atlas (TCGA) were merged based on their corresponding barcodes, and the expression correlation between the paired mRNA and miRNA was tested. Adjusted p-values were calculated using the false discovery rate (FDR) method, and only significant correlations were plotted. Negative correlations were interpreted as potential evidence of negative regulation. The network of miRNA-gene interactions was then constructed using the visNetwork R package.

### 5. Immune association analysis

Immune cell levels within cancers were analyzed using the TCGA data set. A tool called ImmuCellAI [12] was used to evaluate the infiltrates of 24 immune cells. To visualize the data, GSVA scores of relevant genes related to cell death were employed. Spearman correlation analysis was utilized to measure the correlation coefficient between the infiltrates of the immune cells and the expression level of the gene set. All P-values were adjusted by FDR.

### 6. Drug sensitivity analysis

The correlation with drug sensitivity was assessed using a cut-off of remarkable significance (p<1e-5). To further evaluate this correlation, the GSCALite [13] was employed to calculate the area under the dose-response curve (AUC) values of the drugs and gene expression profiles of CENPA in different cancer cell lines. Furthermore, the gene expression profiling data of cancer cell lines from the GDSC [40] and the CTRP [14]were integrated to investigate the sensitivity of the small molecules/drugs with the expression of each gene in the gene set using Spearman correlation analysis.

### 7. Statistical analysis

All statistical analyses were conducted using the R software version 4.0.3. Correlation analysis was performed using the Spearman correlation test, while a Cox proportional hazards model was employed to calculate survival risk and hazard ratio (HR). The prognostic value of each variable was estimated through Kaplan-Meier survival curves and compared through log-rank tests. Comparisons between groups were analyzed through T-test or Analysis of Variance (ANOVA) t-test. If not otherwise stated, the rank-sum test was employed for two sets of data, and a P-value of below 0.05 was considered statistically significant.

## Results

### 1. Single nucleotide variation (SNV) of disulfidptosis cell death genes in cancers

In our analysis of single nucleotide variants (SNVs) related to disulfidptosis-related cell death genes in cancer, we identified FLNA and FLNB as the two most mutated genes in cancer (Fig.1A). For instance, we found 82 FLNA SNV samples in 531 UCEC samples and 53 FLNA SNV samples in 468 SKCM samples. Moreover, our SNV landscapes plot of disulfidptosis-related cell death genes in cancer revealed various SNV type distributions across cancer types(Fig.1B). The majority of SNV types were missense-mutations, while the most common SNV classes were C>T and C>A(Fig.1C). Lastly, we also analyzed and reported the association of SNV of disulfidptosis cell death genes and survival, which revealed that SNVs of most of these genes were not associated with survival (Fig.1D).

**Figure 1.**
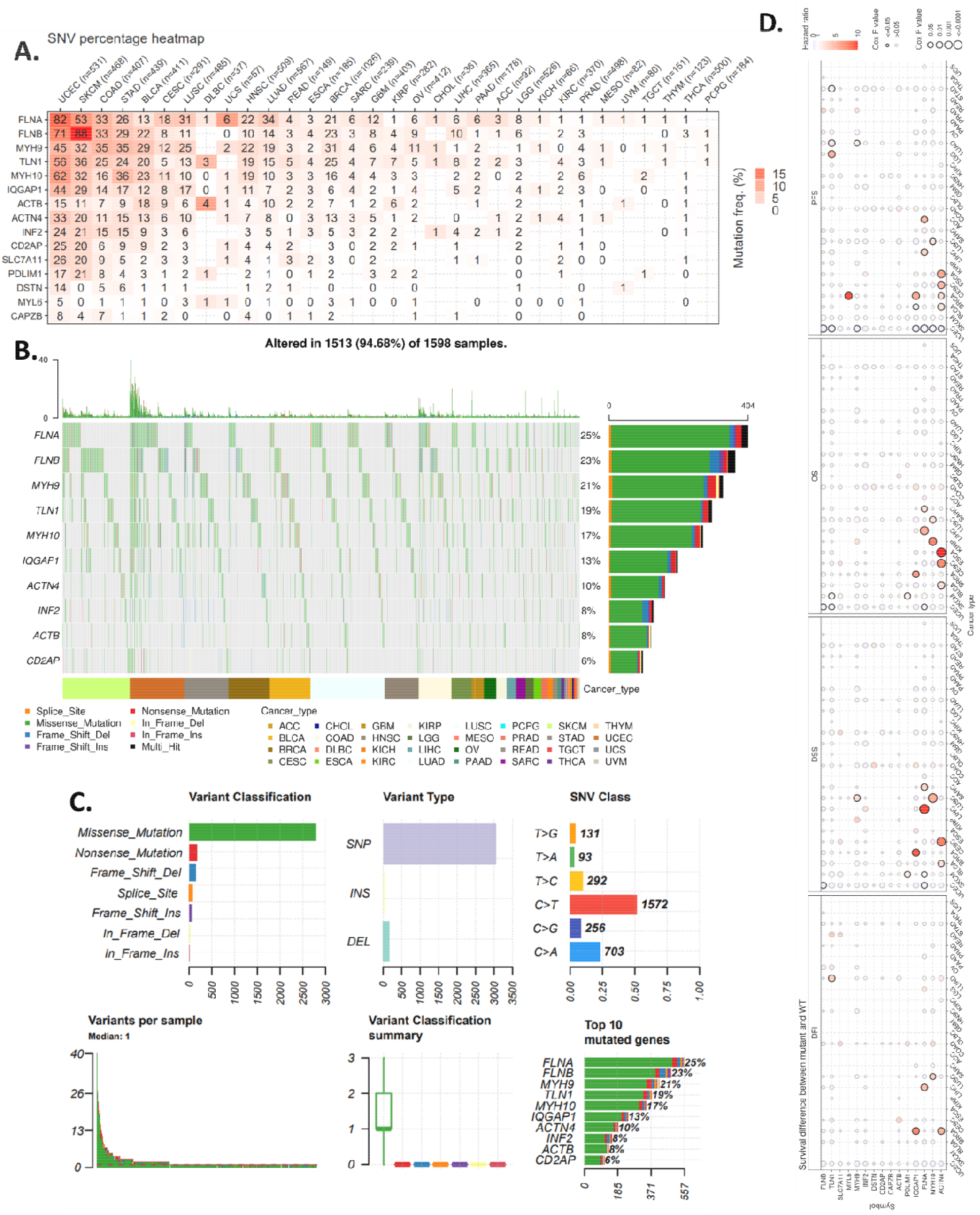
Analysis of single nucleotide variations (SNVs) in disulfidptosis-related cell death genes in cancer. **(A)** Heatmap depicting the mutation frequencies for each gene within the samples, with numbers representing the number of samples with mutated versions of the gene, “0” indicating no mutation in the gene’s coding region, and blank entries indicating a lack of mutation in any region of the gene. Colors represent the mutation frequency. **(B)** SNV landscape plot of the top 10 disulfidptosis-related cell death genes in cancer. **(C)** Summary of SNV classes for disulfidptosis-related cell death genes in cancer, including the count of deleterious mutations, the count of variant types (SNP, INS, and DEL), the count of each SNV class, the count of variants in each sample (a bar represents a sample, the color of the bar corresponds to the color of the variant classification), box plot illustrating the distribution of the count of each variant classification in the sample set (the color of the box corresponds to the color of the variant classification), and the count and percentage of variants in the top 10 mutated genes. **(D)** Survival differences between mutant and wild-type disulfidptosis-related cell death genes.

### 2. Expression profile of disulfidptosis cell death genes in cancer

Our analysis of the expression profiles of disulfidptosis cell death genes in cancer revealed differential expression across multiple cancer types, though the degree of overexpression or underexpression varied (Fig.2A). For instance, PDLIM1 was overexpressed in KIRC but under-expressed in BRCA, LUAD, and PRAD, demonstrating the potential of varying mechanisms of disulfidptosis cell death in different cancer types. Subtype analysis of gene expression revealed BRCA and KIRC as the most significant cancer types (Fig.2B), while a comparison of pathological stage differences indicated KIRC and BLAC as the most notable cancer types (Fig.2C). Additionally, the association of disulfidptosis-related cell death genes with disease-free interval, disease-specific survival, overall survival, and progress-free survival was analyzed, with MESO and LGG as the top cancer types with survival associated with disulfidptosis cell death genes (Fig.2D). Taken together, these results suggest a potential role of regulated expression of disulfidptosis cell death genes in the tumorigenesis of some cancer types.

**Figure 2.**
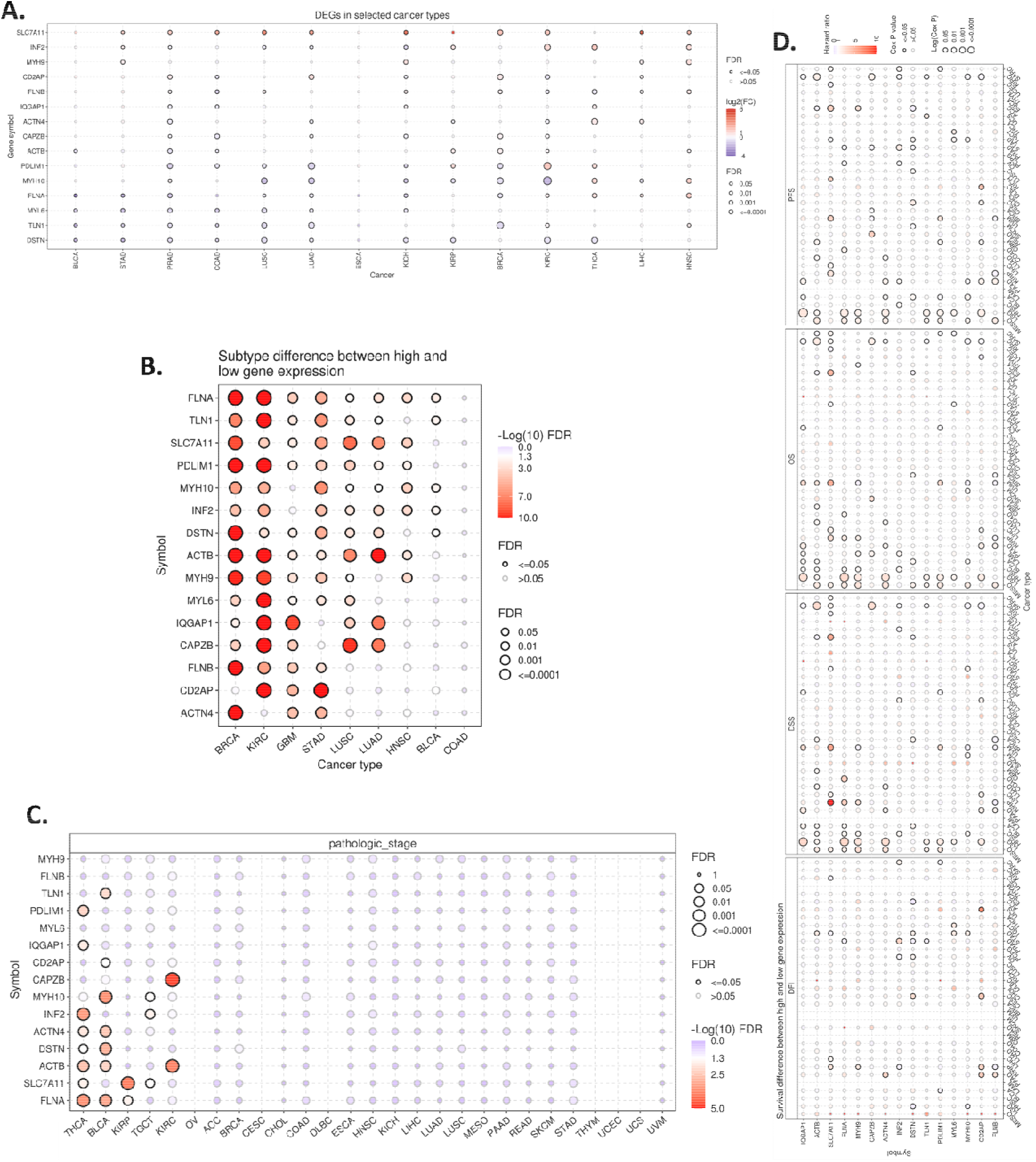
Expression and survival analysis of disulfidptosis-related cell death genes. **(A)** A comparison of mRNA levels between normal and tumor samples. **(B)** Differences in gene expression levels between high and low subtypes. **(C)** Variations in gene expression levels between different pathological stages. **(D)** Survival analysis of disulfidptosis-related cell death genes. The dot size indicates the significance of the gene′s effect on survival in each cancer type, and the color signifies the hazard ratio. DFI: disease-free interval; DSS: disease-specific survival; OS: overall survival; PFS: progress-free survival.

### 3. Copy number variation (CNV) analysis of disulfidptosis-related cell death genes

Our analysis of the copy number variation (CNV) of the disulfidptosis-related cell death genes in cancers revealed that different cancer types had a variety of CNV patterns, with the main CNV types being heterozygous amplification and deletion (Fig.3A). We further plotted the heterozygous amplification and deletion of the disulfidptosis-related cell death genes in cancers (Fig.3B), and calculated the correlation between CNV and expression to evaluate the effect of CNV on expression. This analysis revealed that more than 70% of the cancer types had significant correlations and the top five correlated cancer types were BRCA, LUSC, OV, HNSC, LGG, and LUAD (Fig.3C). Additionally, our analysis of the correlation between CNV and survival across multiple cancer types found that UCEC, KIRP, LGG, and KIRC were the top cancer types where the CNV level was associated with survival (Fig.3D). These results suggest that CNV of the disulfidptosis-related cell death genes directly impacts their expression in cancers and is associated with survival.

**Figure 3.**
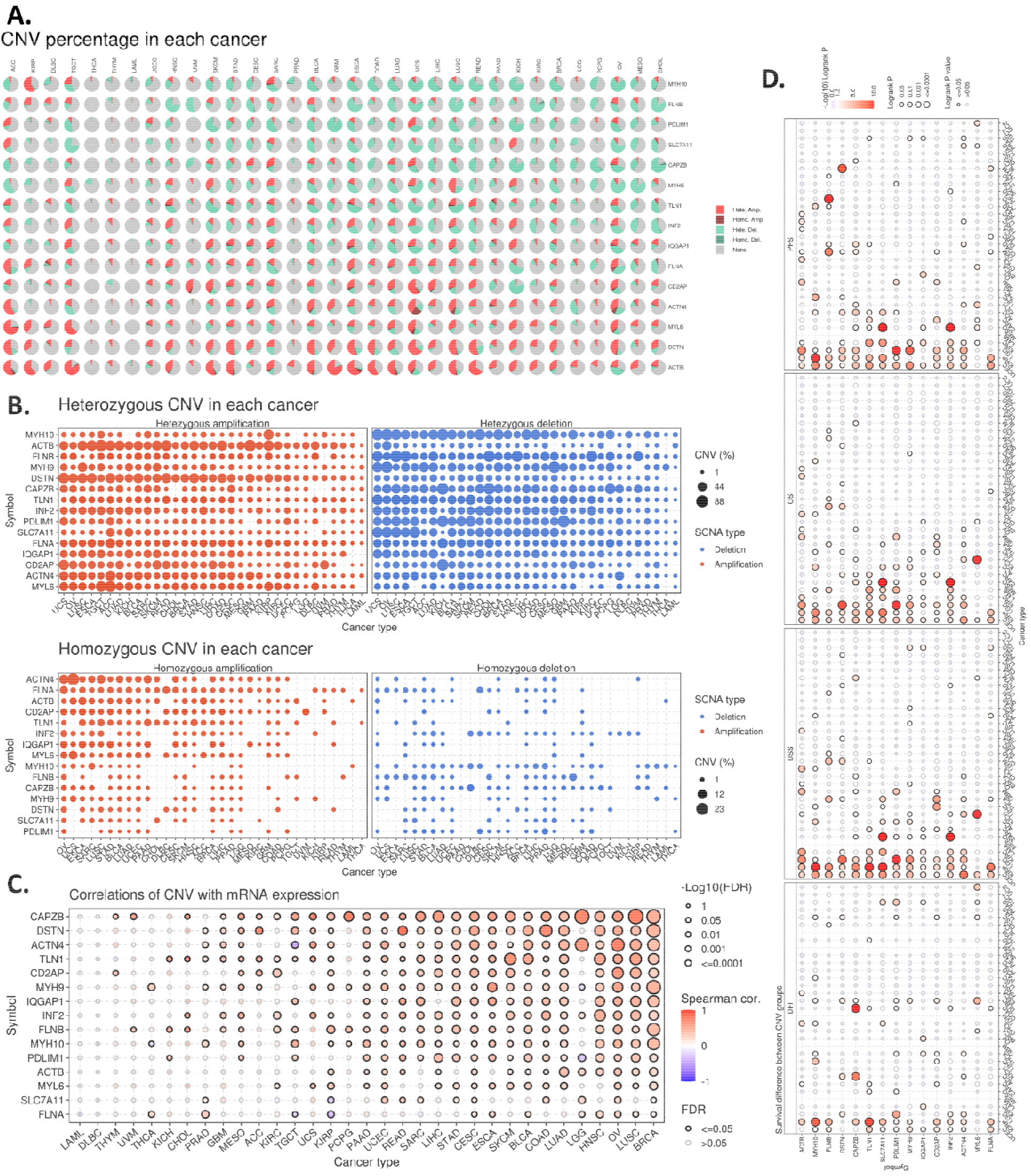
Copy number variation (CNV) analysis of disulfidptosis-related cell death genes in cancers is illustrated. **(A)** Pie chart showing the distribution of CNV across cancers, including Heterozygous Amplification (Hete Amp), Heterozygous Deletion (Hete Del), Homozygous Amplification (Homo Amp), Homozygous Deletion (Homo Del), and No CNV (None). **(B)** Profile of heterozygous CNV showing the percentage of amplification and deletion of heterozygous CNVs for each gene in each cancer. Only genes with >5% CNV in a given cancer are shown. **(C)** Correlation between CNV and mRNA expression. **(D)** Difference in survival between CNV groups.

### 4. Methylation of disulfidptosis cell death genes in cancers

The results of our analysis of methylation showed that, in comparison to normal tissues, the majority of genes had higher methylation levels in tumors in the four most different cancer types (KIRC, THCA, PRAD, and LUSC) (Fig.4A). More genes have higher methylation in tumor than in normal, but some genes in some cancer types have much lower methylation in tumor than in normal such as PDLIM1 in KIRC. Additionally, there was a negative correlation between expression and methylation for most of the genes (Fig.4B). Lastly, there was only a slight correlation between methylation levels and survival of cancer in LGG(Fig.4C).

**Figure 4.**
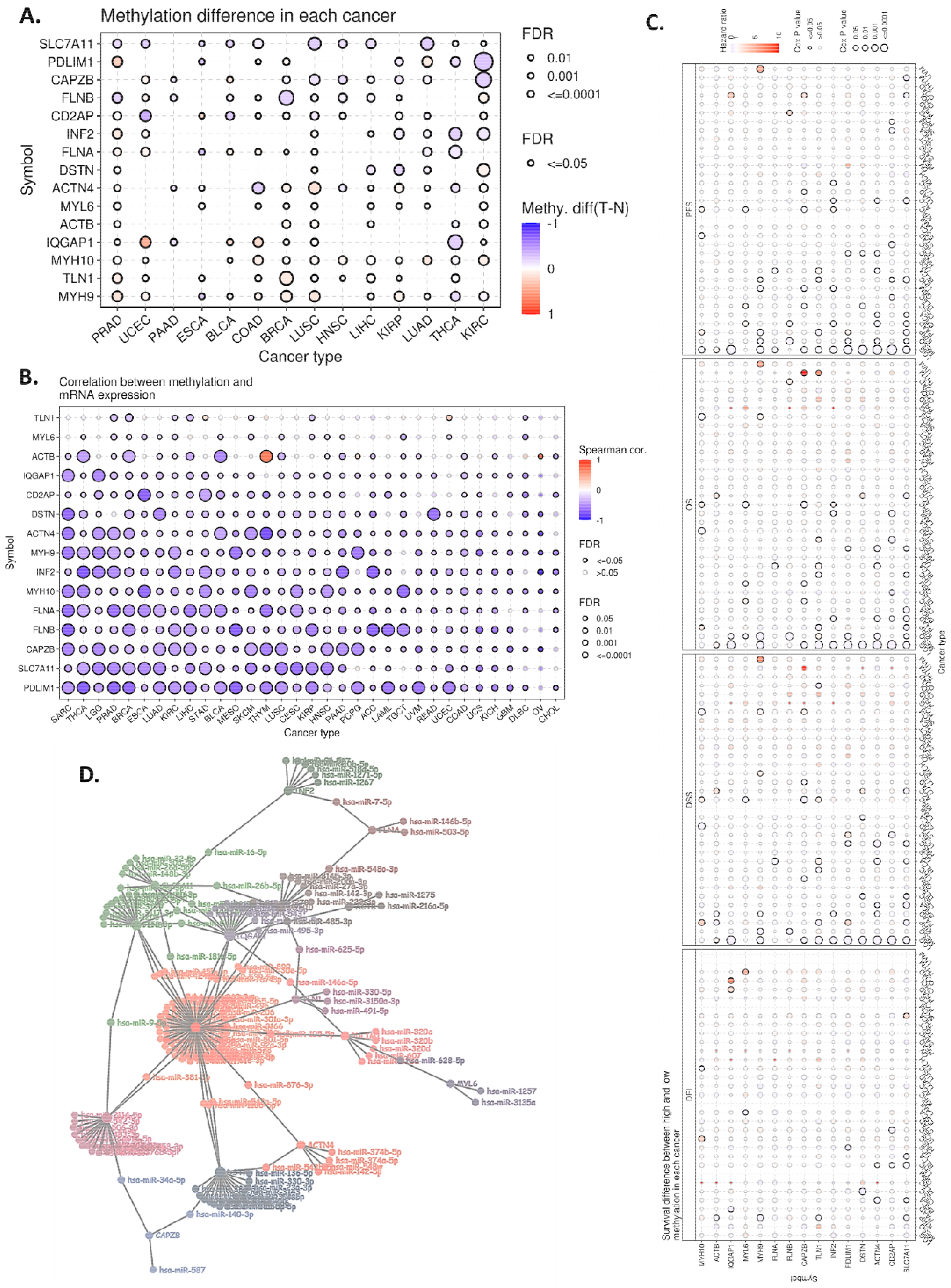
Analysis of methylation patterns of disulfidptosis-related cell death genes in cancer tissues. **(A)** Comparison of methylation profiles between tumor and normal tissue samples. **(B)** Correlation between gene methylation and mRNA expression levels. **(C)** Prognostic significance of methylation differences of disulfidptosis-related cell death genes, with respect to patient survival outcomes. **(D)** A miRNA regulation network of disulfidptosis cell death genes in cancers

### 5. miRNA regulation network of disulfidptosis cell death genes in cancers

The results of our investigation into the potential regulatory effects of miRNA on disulfidptosis-related cell death genes in cancers indicate that there may be a miRNA-mediated regulatory network that controls the expression of these genes. By exploring multiple miRNA databases and analyzing the correlation between target genes and miRNAs, we have been able to construct a network that provides an overview of the potential disulfidptosis-related cell death-regulating miRNAs. This network will be valuable for further research into the regulation of disulfidptosis-related cell death genes (Fig.4D).

### 6. Association profile of disulfidptosis cell death genes and cancer biological processes in cancers

Analysis of the network of pathways related to disulfidptosis-related cell death proteins revealed potential involvement of these proteins in other cancer-related signaling pathways. This suggests a potential association between disulfidptosis cell death and other cancer biological processes. Strikingly, half of the disulfidptosis-related cell death proteins were associated with activation of EMT indicating that the disulfidptosis might associated with cancer migration and invasion (Fig.5).

**Figure 5.**
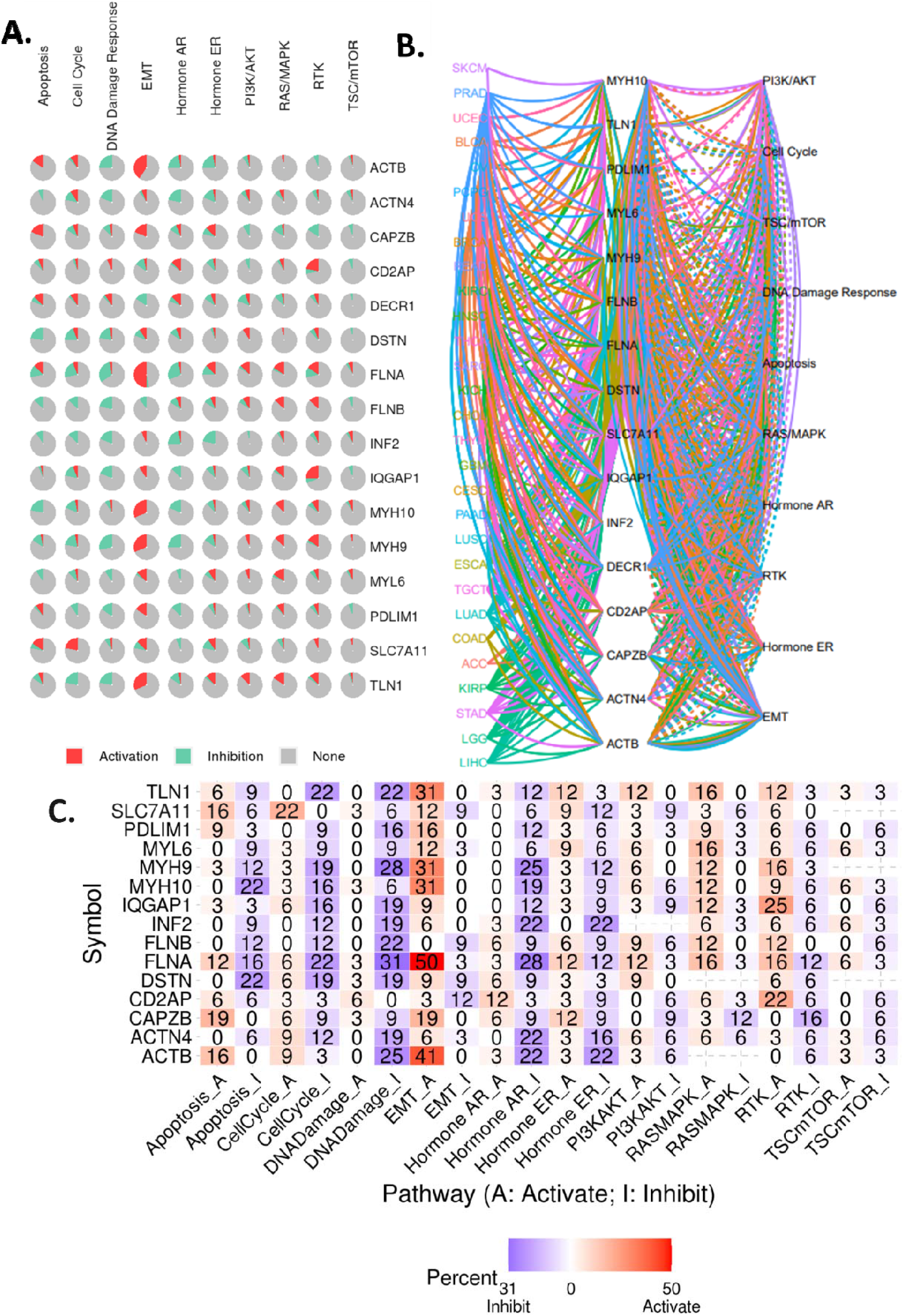
Functional pathway association of disulfidptosis-related cell death genes in cancers. **(A)** The percentage pie chart illustrates the effect of disulfidptosis-related cell death genes on pathway activity. **(B)** A pathway network of disulfidptosis-related cell death genes in cancers. **(C)** The combined percentage of the effect of these genes on pathway activity is also shown.

### 7. Immune and drug sensitivity association profile of disulfidptosis cell death genes in cancers

This study also explored the association of disulfidptosis cell death genes and tumor immune microenvironment. A gene set signature (GSVA score) was constructed to analyze the infiltration levels of multiple immune cells across multiple cancer types. The results showed that the GSVA score was significantly correlated with the infiltration levels of multiple immune cells, particularly gamma delta and B cell (Fig.6A). Furthermore, the correlation between genes and the sensitivity of cell lines to drug compounds was analyzed using GDSC and CTRP databases. The results indicated that the disulfidptosis-related cell death genes were significantly correlated with the sensitivity of cancer cells to multiple compound drugs (Fig.6B). These findings suggest that disulfidptosis-related cell death could affect immune cell infiltration levels in cancers and identified these compounds as potential drugs that might affected by disulfidptosis-related cell death for future study.

**Figure 6.**
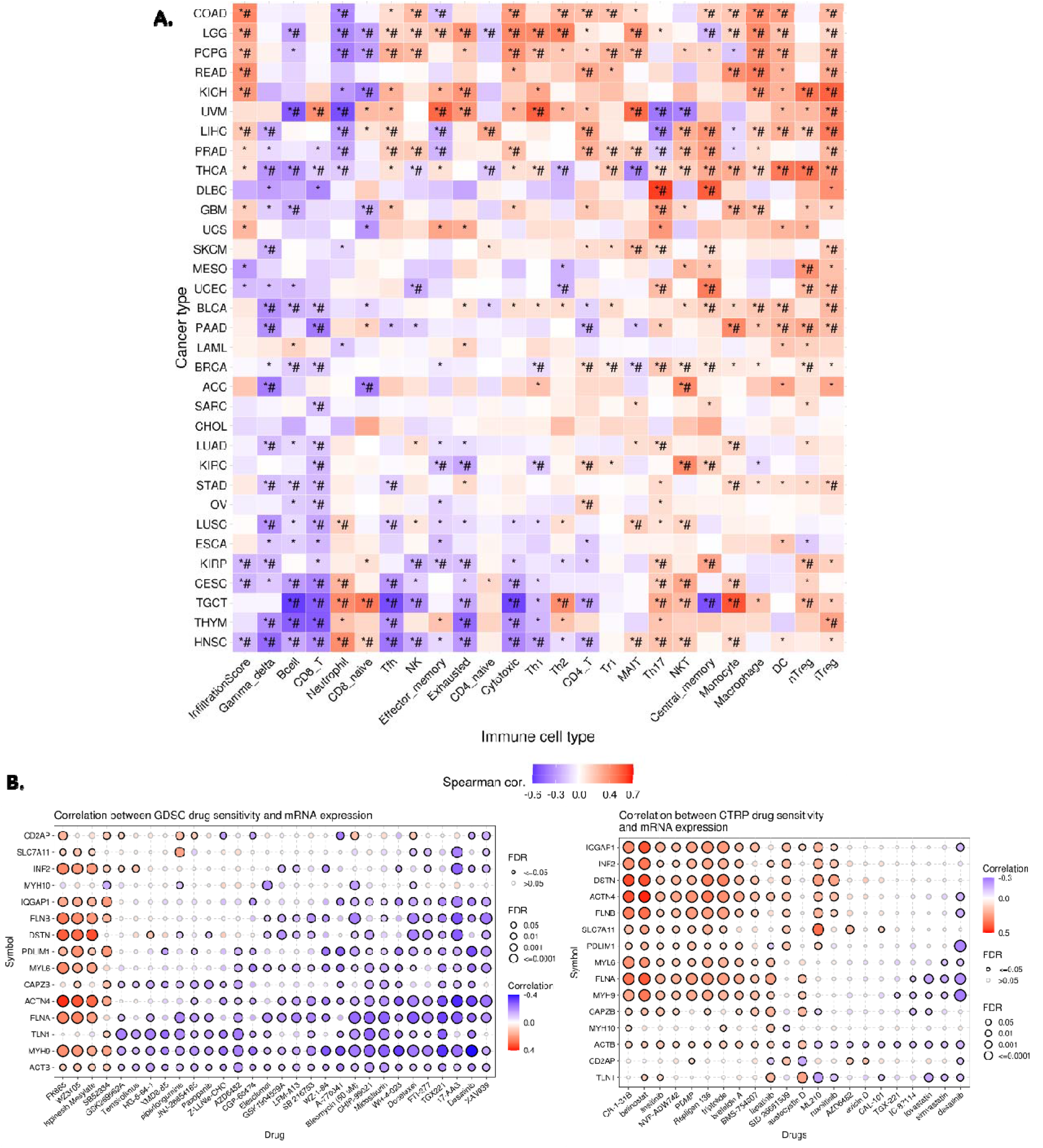
Analysis of disulfidptosis-related cell death genes in cancer types. **(A)** Correlation between GSVA scores of disulfidptosis-related cell death genes and immune cell infiltration levels across cancer types, with statistical significance indicated by *p<0.05 and *#p<0.01. **(B)** Correlation between disulfidptosis-related cell death gene expression and small molecule/drug sensitivity of cancer cell lines, using GDSC and CTRP data. Spearman correlation analysis was used to investigate the relationship between gene expression and small molecule/drug sensitivity (area under IC50 curve).

## Discussion

Cancer is a complex and heterogeneous disease, and its management is a major challenge faced by healthcare professionals worldwide. To effectively combat this disease, a deep understanding of the underlying mechanisms of tumorigenesis is required. Recent studies have identified a novel form of cell death, termed disulfidptosis, which has been shown to be involved in cancer-related processes[2-4,15]. In order to further explore the potential of disulfidptosis as a therapeutic target for cancer treatments, a comprehensive and systematic characterization of the disulfidptosis cell death genes across more than 9000 samples of over 30 types of cancer was conducted. By mining multi-omics profiling data, the expression and regulation of disulfidptosis cell death genes in cancer contexts were elucidated. Moreover, the potential associations between disulfidptosis cell death and other common cancer pathways were analyzed, providing an overall picture of disulfidptosis-related cell death in cancer. The findings of this study may open up potential new avenues for the development of novel cancer treatments.

This study explored the disulfidptosis pathway and its potential therapeutic targets for cancer treatment. Disulfidptosis is a newly discovered pathway, which involves the regulated formation of reactive disulfide bonds in proteins and non-protein molecules. The downstream genes of the disulfidptosis pathway were not investigated in this study; however, the disulfidptosis-based target therapy might largely depend on the downstream elements. Thus, further study of these downstream targets will be valuable for the development of disulfidptosis-based target therapy. In addition, the miRNA network analysis also provided potential therapeutic targets for disulfidptosis-based therapy. The interactions were provided by the databases including some experimentally verified data and predicted data. Some of these interactions were validated by experimental evidence; however, further studies in this field are still needed. This study provided potential disulfidptosis-related miRNA for future research, with the aim of developing effective cancer therapies.

The disulfidptosis pathway is a double-edged sword with implications for both normal cellular homeostasis and cancer treatment. Variations in the expression of disulfidptosis genes can lead to lethal disorders in humans, yet, the same pathway may also be used to aid in cancer treatment. Our evidence has suggested that the disulfidptosis pathway may be associated with the infiltration of multiple immune cells across different cancer types. However, it is still not clear whether disulfidptosis functionally affects immune cells or whether immune cell infiltration can induce disulfidptosis. In addition, the disulfidptosis core gene signature has been proposed as a useful predictor for the response to immune therapy and survival of patients. However, further validations with prospective evidence are needed to confirm its efficacy. As cancer is a complex and heterogeneous disease, understanding the role of disulfidptosis in cancer is essential for optimizing the efficacy of cancer treatments.

The association between disulfidptosis-related cell death and drug sensitivity is still not clear. However, our analysis revealed that the expression of disulfidptosis cell death genes in cancer is associated with the sensitivities of cancer cells to multiple chemicals. Multiple cancer cell lines were integrated to provide a general view that disulfidptosis cell death might potentially affect drug resistance. Additionally, SLC31A1 was also reported to be associated with drug resistance [5,16,17]. Although the patterns of the correlations were different, including some positive and some negative correlations, this indicated that the mechanisms of disulfidptosis-related cell death in drug resistance might be different among different drugs. Thus, further detailed studies are required to explore the mechanisms of disulfidptosis-related cell death in different drug resistance.

Although the analysis results obtained from studying the role of lipoylated proteins in disulfidptosis-related pathways are intriguing, they are sometimes contrary to what was anticipated in terms of their impact on cell death. This is understandable, as there is no established consensus on the mechanism by which the aggregation of lipoylated proteins leads to cell death in disulfidptosis-related pathways. Consequently, it is unclear whether higher or lower expression of these proteins is more detrimental or beneficial for cell death. This study consequently aims to investigate the effects of cystine transporter solute carrier family 7 member 11 (SLC7A11) expression on cell death in cancer cells. It is hypothesized that the accumulation of intracellular disulfide, such as cystine, can be highly toxic for cells and deplete the reduced form of nicotinamide adenine dinucleotide phosphate (NADPH) pool, resulting in a rapid build-up of disulfide molecules and consequent cell death when coupled with glucose starvation. As cancer is a multifaceted disease characterized by abnormal cell growth that can spread to other parts of the body, understanding the underlying mechanisms of disulfidptosis-induced cell death is of paramount importance for the development of novel therapeutic strategies to combat this disease and others. The findings of this study may provide insight into the role of disulfidptosis as a novel form of cell death in cancers, as well as potential therapeutics to target it.

## Conclusion

This study aimed to evaluate the potential relevance of disulfidptosis-related cell death genes in cancer. Analyses of cancer types revealed that disulfidptosis-related genes can be aberrantly expressed, have an altered level of mutation, and are linked to cancer survival and cross-pathway interactions. Furthermore, the implications of these genes on the cancer microenvironment and drug resistance suggest their potential value in cancer therapies. As such, disulfidptosis-related genes may be developed as suitable biomarkers for cancer diagnosis, prognosis, and therapeutic applications.

## Supporting information

miRNA network

## List of abbriviations

ACC: Adrenocortical carcinoma
BLCA: Bladder Urothelial Carcinoma
BRCA: Breast invasive carcinoma
CESC: Cervical squamous cell carcinoma and endocervical adenocarcinoma
CHOL: Cholangio carcinoma
COAD: Colon adenocarcinoma
DLBC: Lymphoid Neoplasm Diffuse Large B-cell Lymphoma
ESCA: Esophageal carcinoma
GBM: Glioblastoma multiforme
HNSC: Head and Neck squamous cell carcinoma
KICH: Kidney Chromophobe
KIRC: Kidney renal clear cell carcinoma
KIRP: Kidney renal papillary cell carcinoma
LAML: Acute Myeloid Leukemia
LGG: Brain Lower Grade Glioma
LIHC: Liver hepatocellular carcinoma
LUAD: Lung adenocarcinoma
LUSC: Lung squamous cell carcinoma
MESO: Mesothelioma
OV: Ovarian serous cystadenocarcinoma
PAAD: Pancreatic adenocarcinoma
PCPG: Pheochromocytoma and Paraganglioma
PRAD: Prostate adenocarcinoma
READ: Rectum adenocarcinoma
SARC: Sarcoma
SKCM: Skin Cutaneous Melanoma
STAD: Stomach adenocarcinoma
TGCT: Testicular Germ Cell Tumors
THCA: Thyroid carcinoma
THYM: Thymoma
UCEC: Uterine Corpus Endometrial Carcinoma
UCS: Uterine Carcinosarcoma
UVM: Uveal Melanoma

## Ethical Approval and Consent to participate

Not applicable.

## Consent for publication

Not applicable.

## Availability of supporting data

The authors of this paper sourced the raw data and conducted a raw analysis. The authors are willing to share the data with reasonable requests from a corresponding author.

## Authors′ contributions

All the analysis works were done by Hengrui Liu. Hengrui Liu and Tao Tang wrote the paper. Tao Tang edited the paper and supervised the project.

## Acknowledgements

The author thanks the support of the GSCALite[13], Dr. Jieling Weng, Weifen Chen, Zongxiong Liu, and Yaqi Yang.

## Authors′ information

Hengrui Liu is a Principal Investigator in Tianjin Yinuo Biomedical Co., Ltd. Tao Tang is cancer molecular diagnostic technician at Sun Yat-Sen University Cancer Center.

